# Revealing an extended critical window of recovery post-stroke

**DOI:** 10.1101/458745

**Authors:** Belen Rubio Ballester, Armin Duff, Martina Maier, Monica Cameirao, Sergi Bermudez, Esther Duarte, Ampar Cuxart, Susana Rodríguez, Paul F.M.J. Verschure

## Abstract

The impact of rehabilitation on post-stroke motor recovery and its dependency on the patient’s chronicity remain unclear. The existence and regularity of a, so called, proportional recovery rule across a range of functional deficits and therapies supports the notion that functional interventions have little or no impact beyond spontaneous recovery rates in a ‘critical window of recovery’ which lasts from 3 to 6 months post-stroke. In this meta-analysis, we apply a bootstrap analysis method to assess the overall impact of a specific VR-based rehabilitation protocol for the upper extremities on a homogeneous sample of 219 individuals with hemiparesis at various stages post stroke. Our analysis uncovers a precise gradient of sensitivity to treatment that expands more than one year beyond the limits of the so-called ‘critical window of recovery’. These findings redefine the limits of the so-called ‘critical window of recovery’ and suggest that stroke-derived plasticity mechanisms do facilitate functional recovery even at the chronic and late chronic stage.

## 1 INTRODUCTION

The absolute incidence of stroke will continue to rise globally with a predicted 12 million stroke deaths in 2030 and 60 million stroke survivors worldwide (Feigin et al., 2014). Stroke leads to focal lesions in the brain due to cell death following hypoxia and inflammation, affecting both gray and white matter tracts (Corbetta et al., 2015). After a stroke, a wide range of deficits can occur with varying onset latencies such as hemiparesis, abnormal posture, spatial hemineglect, aphasia, and spasticity, along with affective and cognitive deficits, chronic pain and depression (Teasell et al., 2015). Due to improved treatment procedures during the acute stage of stroke (e.g. thrombolysis and thrombectomy), the associated reduction in stroke mortality has led to a greater proportion of patients facing impairments and needing long-term care and rehabilitation. However, prevention, diagnostics, rehabilitation and prognostics of stroke recovery have not kept pace (Veerbeek et al., 2014).

Post-stroke recovery largely follows a non-linear logarithmic trajectory (Kwakkel et al., 2004). On this basis, the main goal of rehabilitation is to affect the slope (i.e. faster recovery) and the asymptote (i.e. the level of functioning of the stroke patient) of this recovery curve. A variety of rehabilitation approaches are available such as occupational therapy, constrained induced movement therapy, physical movement therapy, virtual reality-based therapies, and/or neuromuscular stimulation however most of these therapies have not shown an impact beyond what is expected from spontaneous recovery (Krakauer and Cort, 2018 Winstein et al., 2016). Post-stroke mechanisms that promote recovery include revascularization, axonal sprouting, neurogenesis and rewiring and the rebalancing of excitation and inhibition in cortical networks which can be misinterpreted as the effects of behavioural therapies. Indeed, animal studies suggest the existence of a specific time window of increased responsiveness to training, the so-called ‘critical window of recovery’, which may be associated to the aforementioned recovery mechanisms (Biernaskie, 2004 Murphy and Corbett, 2009). However, a similar temporal structure of recovery has not been validated in humans yet and the optimal timing for treatment thus remains unclear. Three ongoing clinical trials are exploring this specific question: McDonnell et al. (McDonnell et al., 2015), the SMARTS 2 trial (NCT02292251)(Krakauer and Cort, 2018), and the CPASS trial (Dromerick et al., 2015). Here, we directly address this critical question in post stroke rehabilitation by performing an analysis across a series of clinical studies that have all deployed the same VR based rehabilitation protocol at various stages post stroke from the acute to the late chronic stage.

In order to understand the relationship between rehabilitation and chronicity, we perform an analysis of longitudinal clinical data from 219 stroke patients who followed the same VR-based training protocol which targets the rehabilitation of upper extremity (UE) functionality using the Rehabilitation Gaming System (RGS) (Cameirão et al., 2010). The RGS is a VR-based framework for rehabilitation in which arm and finger movements are displayed on a screen in a first-person perspective, realizing a paradigm that combines goal-oriented action execution, motor imagery, and action observation (Figure S1). To analyze the impact of RGS across the poststroke continuum of care, we propose a new methodology for the exploration of stroke recovery that addresses the problem of the high inter-subject variability. We show that the intervention we analyze has a significant impact on the function of the UE at all periods post stroke considered, uncovering a gradient of enhanced recovery and high sensitivity to the intervention that extends beyond 18 months post-stroke.

## 2 MATERIALS AND METHODS

### 2.1 The mechanisms and principles of the Rehabilitation Gaming System (RGS)

This work builds on previous research conducted with the RGS, a VR-based tool that promotes functional recovery post-stroke through goal-oriented embodied sensorimotor stimulation. A number of studies suggest the effectivity of RGS protocols for overcoming upper limb motor deficits (Cameirão et al., 2012 da Silva Cameirão et al., 2011; Ballester et al., 2016, 2015, 2017). These protocols rest on principles that are derived from the Distributed Adaptive Control theory of mind and brain (Verschure, 2012), which places recovery in the context of the acquisition and expression of goal-oriented voluntary behavior driven by perception, memory, value and goals and the optimization of perceptual and behavioral prediction. Through these mechanisms, RGS aims at promoting neuronal and functional reorganization by engaging the affected area of the brain through non-invasive exposure to multisensory stimulation (Prochnow et al. 2013). Specifically, RGS combines goal-oriented action execution with a first-person observation of the corresponding movement in VR (Figure S1), in addition to that activating undamaged primary and secondary motor areas, and promoting functional reorganization. Indeed, it has been shown that the observation of hand movements leads to an increase in cortical excitability when the orientation of the observed upper limbs and the point of view of the observer are congruent (Maeda et al., 2002). The principle of embodied ideomotor training is integrated into a range of goal-oriented tasks, exercising different aspects of motor and cognitive function that have been shown to enhance retention of new motor schema inducing more substantial treatment effects (Timmermans et al., 2010). RGS individualizes training, where each trial of each session is customized to the capabilities of the user and the specific training objectives using machine-learning techniques (Nirme et al., 2011). The task considered here, Spheroids, consists in reaching, grasping, and releasing spheres into color-matched boxes (Cameirão et al., 2010). The task is divided into 3 subtasks that are structured in time, and progress from proximal to distal movements. This design introduces task variability during the training sessions and provides a practice schedule that is structured, including rest periods. These components have shown to optimize the acquisition, retention, and generalization of motor skills (Yamazaki et al., 2015; Hanlon, 1996). In order to promote the usage of the affected limb, the scenarios contain contextual restrictions that limit both the overuse of the non-affected arm and compensatory movements, thus supporting use-dependent restoration towards non-pathological motor patterns. Moreover, the patient and the therapist receive explicit feedback, including both information about performance and results, thus reinforcing the execution of appropriate successful goal-oriented movements and maximizing long-term retention of motor memories (Abe et al., 2011). Overall, the RGS training protocols integrate five main principles for motor recovery: 1) include self-paced individualized intense practice, in ecologically valid settings, 2) limit overcompensation, 3) promote goal-oriented tasks that are structured in time, 4) facilitate motor imagery though embodied training, and 5) provide multimodal feedback.

### 2.2 Identification of studies

To evaluate the effectivity of RGS in various stages post-stroke, we first identify all published longitudinal studies that tested the effectivity of the Spheroids scenario of the RGS for the recovery of the upper extremity motor function. Studies were excluded if they were not longitudinal, included less than five subjects per group, did not include the standard RGS Spheroids protocol, did not target upper extremities, or did not provide information on standard clinical scales at baseline, end of the treatment and follow-up. In all the selected studies, recruited patients met the following inclusion criteria: 1) ischemic strokes (Middle cerebral artery territory) or hemorrhagic strokes (intracerebral), 2) mild-to-moderate upper limb hemiparesis (Medical Research Council scale for proximal muscles > 2) after a first-ever stroke, 3) age between 45 and 85 years old, 4) the absence of any significant cognitive impairment (Mini-Mental State Evaluation > 22). Overall, the ten studies selected were divided into seventeen conditions depending on the specific characteristics of the patients and the requirements of the treatment provided (Table 1 including the patients’ chronicity, type of intervention, number of patients tested, age, days post-stroke, lateralization, sex and stroke type). Conditions 2, 3, 7 and 12 combine RGS-based training with supervised OT. Study 13 tested the application of RGS in a domiciliary setting. A formal risk of bias analysis on the primary outcome (i.e. change in the UE-FM) was performed using ROBINS-I tool (Supporting Information, Table S1), covering the evaluation of confounding variables, recruitment for participants, intervention classification, deviations from intended interventions, missing data, measurement of outcomes and selection of reported results.

### 2.3 Outcome Measures

The primary outcome considered in our study was the UE motor function at the end of therapy, as measured by two standardized clinical scales (i.e. UE-FM and CAHAI scales). Since previous studies have shown that the UE-FM shows excellent reliability, responsiveness and validity properties (Wei et al., 2011), the use of this scale was prioritized. Secondly, we analyzed improvement measures captured by the CAHAI (Barreca et al., 2005), which evaluates the UE bilateral function in the performance of specific iADLs. Score changes in the UE-FM and CAHAI were used as measures of structural and functional motor recovery respectively.

### 2.4 Statistical Analysis

We performed two analyses. Firstly, we reported primary outcome measures on the FM-UE and CAHAI in absolute terms. In this case, we quantified improvement using mean differences and 95% CIs. For all analyses, statistical significance levels were set at p < 0.05. Secondly, to analyze the overall effect of RGS, we applied a bootstrap method (Efron and Tibshirani, 1986) to compare outcome measures from 178 patients performing RGS-based training and 41 control subjects following conventional rehabilitation protocols. This method allowed us to bundle the data of the different studies into two intervention groups obtaining accurate estimates of inter-group variability. In this way, we overcame the risk of bias across studies due to the variation in treatment intensity and response rates among experimental and control groups, which is a common limitation in the meta-analysis of clinical trials (Makuch and Johnson, 1989). This homogenized data was generated by separating improvement measures at the different time intervals (end of treatment and follow-up) of the studies and allocating them to either being an RGS condition, OT condition in which patients followed standard protocols and follow-up (i.e. no-therapy). We then calculated the improvement rate per week-normalized within-subjects according to their respective recovery potential (i.e., the improvement observed normalized to the total amount that each can gain concerning their baseline) in standardized clinical scales. The normalized improvement on scale *i* at time *t*, was defined as:

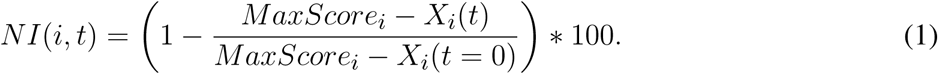

**Table 1.** Overview of the RGS studies included in the analysis.

| ID | Group | Stage Post-Stroke | Intervention | N | Age Mean(SD) | TSO (days) | RUE (%right) | Sex (%male) | Oxf.Class. | Reference |
| --- | --- | --- | --- | --- | --- | --- | --- | --- | --- | --- |
| 1 | Control | Acute | 12w;5d/w;20min; | 5 | 69(19) | 12(10) | 40 | 80 | 2/1/0/1/1 | da Silva Cameirão et al., 2011; Duff et al., 2013 |
| 2 | RGS | Acute | 3w;5d/w;20min; | 5 | 70(22) | 13(3) | 60 | 20 | 1/2/0/1/1 | Rodriguez et al., 2011; Duff et al., 2013 |
| 3 | RGS | Acute | 12w;3d/w;20min; | 10 | 63.5(29) | 11(17) | 40 | 30 | 2/2/2/3/1 | da Silva Cameirão et al., 2011; Duff et al., 2013 |
| 4 | Control | Acute | 3w;5d/w;20min; | 5 | 64(16) | 13(5) | 60 | 60 | 2/2/0/0/1 | Rodriguez et al., 2011; Duff et al., 2013 |
| 5 | Control | Acute | 12w;5d/w;20min;NSG | 4 | 65(28) | 13.5(12) | 75 | 50 | 1/0/1/1/1 | da Silva Cameirão et al., 2011 |
| 6 | Control | Acute | 12w;5d/w;20min;IOT | 5 | 56(27) | 15(11) | 40 | 40 | 1/1/2/0/1 | da Silva Cameirão et al., 2011 |
| 7 | RGS | Subacute | 3w;3d/w;20min; | 49 | 61(43) | 70(375) | 30.6 | 30.6 | 11/9/13/1/15 | Unpublished |
| 8 | Control | Subacute | 3w;5d/w;20min | 4 | 57(17) | 90(226) | 0 | 50 | 4/0/0/0/0 | Unpublished |
| 9 | RGS | Chronic | 6w;5d/w;30min;+AM | 9 | 63(31) | 400(5805) | 33.3 | 33.3 | - | Ballester et al., 2016 |
| 10 | RGS | Chronic | 6w;5d/w;30min | 9 | 57(36) | 735(4471) | 11.1 | 55.6 | - | Ballester et al., 2016 |
| 11 | Control | Chronic | 3w;5d/w;20min;domiciliary | 18 | 68.5(40) | 751(1536) | 33.3 | 50 | 6/2/4/0/6 | Ballester et al., 2017; Nirmé et al., 2013 |
| 12 | RGS | Chronic | 3w;5d/w;20min;+OT | 20 | 64.5(37) | 770(2789) | 40 | 30 | 7/0/4/0/9 | Cameirão et al., 2012 |
| 13 | RGS | Late Chronic | 3w;5d/w;20min;domiciliary | 17 | 61.5(43) | 997(2987) | 47.1 | 35.3 | 4/3/4/0/6 | Ballester et al., 2017; Nirmé et al., 2013 |
| 14 | RGS | Late Chronic | 4w;5d/w;30min;haptics | 14 | 63(45) | 1051(3250) | 50 | 57.1 | 6/0/4/1/3 | Cameirão et al., 2012 |
| 15 | RGS | Late Chronic | 3w;5d/w;30min | 15 | 58(60) | 1261(2041) | 33.3 | 46.7 | 0/1/2/0/12 | Unpublished |
| 16 | RGS | Late Chronic | 4w;5d/w;30min | 16 | 69.5(46) | 1536(3891) | 43.8 | 62.5 | 6/4/5/0/1 | Cameirão et al., 2012 |
| 17 | RGS | Late Chronic | 4w;5d/w;30min;exoskeleton | 14 | 60(32) | 1758(2880) | 35.7 | 71.4 | 3/1/4/1/5 | Cameirão et al., 2012 |

Intervention: duration of included protocols indicated per number of weeks (w), days per week (d/w), and minutes (min) of Occupational Therapy (OT) and VR-based therapy per day. N: sample size in the experimental group. TSO: time since stroke (days). NSG: protocol based on non-specific gaming system (i.e., Nintendo Wii). IOT: protocol including intensive occupational therapy. AM: protocol including the amplification of movements in VR. Haptics: protocol including delivery of haptic feedback during training. RUE: percentage of patients with right upper-extremity affected. Sex: percentage of males. Oxf. Class: Count of stroke types (lacunar stroke (LACS) / partial anterior circulation stroke (PACS) / total anterior circulation stroke (TACS) / or posterior circulation stroke (POCS)) according to the Oxford Stroke Classification scale.

where *X_i_*(*t*) is a given measure on scale *i* at time *t*, and *X_i_*(*t* = 0) refers to the corresponding baseline score. This normalization adjusts for the risk of bias due to selecting participants with variable baseline severity of upper limb hemiparesis (Saposnik and Levin, 2011; Medica et al., 2015). We computed an NI value for three-time intervals: pre-, post-assessments, and long-term follow-up. For example, for a patient who followed RGS training with a baseline, end of the treatment, and follow-up assessment, we calculated the NI per week for two periods: baseline to end of the treatment and end of the treatment to long-term follow-up. The measured change from baseline to end of the treatment was allocated to the RGS or the OT group, while the change from the end of the treatment to follow-up was assigned to the follow-up group. If a patient had multiple follow-up assessments at different time points, all the follow-up measures were assigned to the follow-up group. To facilitate the exploration and replication of our findings, we are publishing the complete dataset and analysis source-code as freely available and open, which will be accessible at an OSF repository and available on publication.

## 3 RESULTS

### Impact assessment of individual studies

We identified ten studies testing the RGS Spheroids protocol (see Methods, Figure S1). We excluded four studies for different reasons: two studies were cross-sectional and did not report measures of motor recovery, one study did not report measures of motor recovery using clinical scales, and another study employed a modified version of the Spheroids scenario. A total of six articles including data from 219 stroke survivors assigned to 17 VR-based rehabilitation conditions (Table 1) were selected for the qualitative and the quantitative analysis of results. The methodological homogeneity of the RGS studies allowed us to combine their results and to compare groups of patients in three time windows defined by the time elapsed from the onset of the stroke to the time of baseline evaluation: acute (< 3 weeks), subacute (3 weeks to 6 months), early chronic (6–18 months), and late chronic (> 18 months). We observed significant UE structural gains after treatment, both in acute (median 20.0 ± 7.9 MAD, p < .01, Wilcoxon sign-rank) and subacute patients (median 8.0 ± 5.6 MAD, *p* < .01), as measured by UE-FM (Figure 1 A, Table 2). These gains were accompanied by an improved performance in iADLs in both acute (median 42.5 ± 14.1 MAD *p* < .01) and subacute groups (median 7.0 ± 10.5 MAD, *p* < .01), as measured by CAHAI after treatment (Figure 1 B, Table 2). More interestingly, at the chronic and late chronic stage, the RGS showed overall effectiveness in facilitating improvements in UE-FM (ranging from median 2.7 ± 3.8 MAD to median 7.0 ± 3.6 MAD, *p* < .05) and CAHAI (median 1.0 ± 3.8 MAD to median 8.0 ± 5.6 MAD, *p* < .05). However, the application of the RGS to domiciliary protocols (chronic at home) showed no significant effects in UE-FM but induced statistically significant but non-clinically relevant gains in the execution of iADLs (median 1.0 ± 1.6 MAD, *p* < .01). Surprisingly, a dosage-matched RGS study conducted in the clinic on late chronic patients had an impact on UE-FM (condition 15 in Table 1, median 3.0 ± 4.1 MAD, *p* < .01; Figure 1 A and B, condition Late Chronic 3w). Furthermore, we observed a dependency between the number of days post-stroke before the start of the RGS therapy and the improvements in motor function as measured by UE-FM and CAHAI.

In general, acute patients that trained with the RGS showed a greater recovery when compared to the standard treatment (Supporting Information, Figure S2) and subacute and chronic patients that followed a dosage-matched RGS-based treatment (Figure 1, Table 2). The analysis of follow-up measures illustrates that improvements were retained in all groups. Interestingly, the subacute subgroup training with RGS exhibited a significant improvement during the follow-up period (3 months) both in UE-FM (median 2.0±5.3 MAD, *p* < 0.01) and CAHAI (median 3.0±11.7 MAD, *p* < 0.01) (Supporting Information, Figure S3 A and B). The acute groups, however, showed higher inter-individual variability and non-significant gains from the end of the therapy to the follow-up.

**Figure 1.**
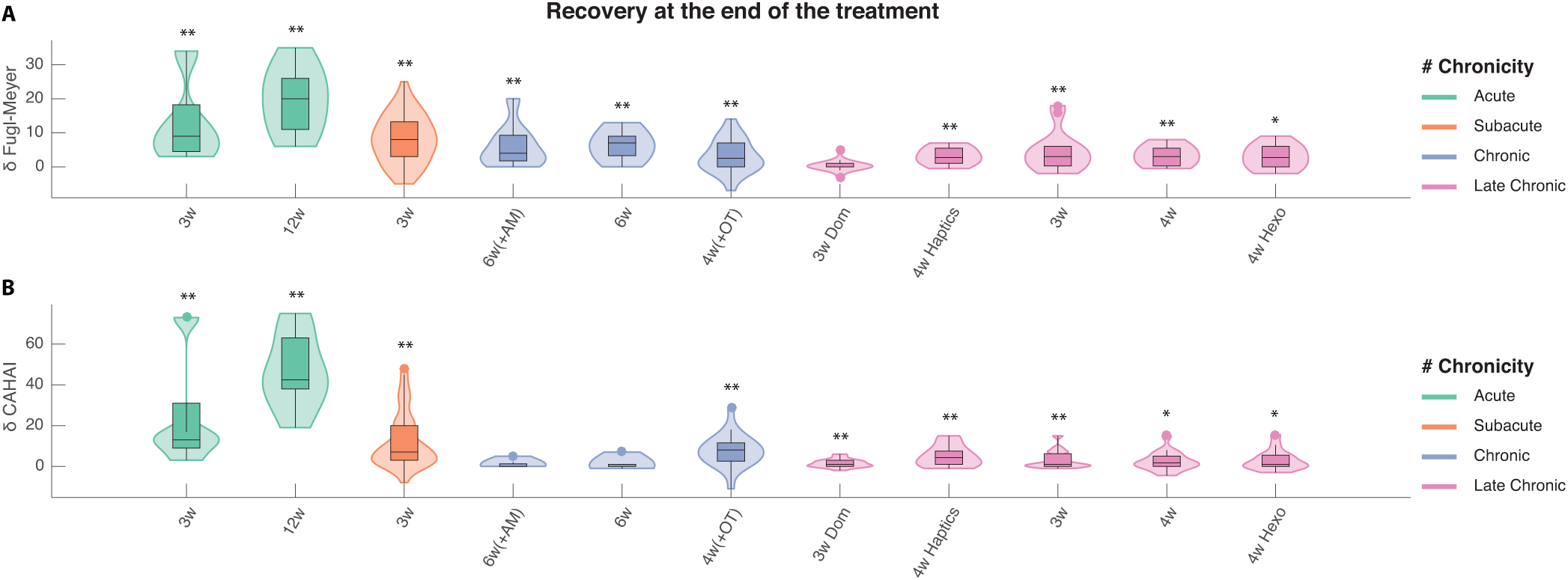
Effect of RGS-based treatment from the start to the end of the therapy. Impact measured on upper limb motor function in the Upper Extremity section of the Fugl-Meyer scale (UE-FM) (top) and performance in iADLs captured by the Chedoke Arm and Hand Activity Inventory (CAHAI) (bottom). The effect represents a change in each scale from the start to the end of the treatment. The length of each intervention is indicated in weeks (w). Notice that the horizontal axis refers to the RGS conditions listed in Table 1 and follow the same order. Shaded areas indicate the data distribution color coded according to the chronicity of stroke patients participating in each study: acute (green), subacute (orange), and early (blue) to late (purple) chronic stage. * for p-value< 0.05 and ** for p-value< 0.01.

**Table 2.** Impact on Recovery at T1 and Follow-up by treatment condition.

| ID | Change from Baseline to T1 |  |  |  | Change from T1 to Follow-Up |  |  |  |
| --- | --- | --- | --- | --- | --- | --- | --- | --- |
| | Median FM ± MAD | Median CAHAI ± MAD | p-value (FM) | p-value (CAHAI) | Median FM ± MAD | Median CAHAI ± MAD | p-value (FM) | p-value (CAHAI) |
| 1 | 24.000 ± 14.240 | 42.000 ± 14.640 | - | - | 8.00 ± 3.60 | 3.00 ± 5.12 | - | - |
| 2 | 9.000 ± 8.560 | 13.000 ± 19.840 | - | - | 6.00 ± 11.04 | 12.00 ± 11.68 | - | - |
| 3 | 20.000 ± 7.900 | 42.500 ± 14.080 | .002 | .002 | 0.00 ± 2.04 | 0.00 ± 3.00 | .516 | > .999 |
| 4 | 18.000 ± 6.240 | 10.000 ± 13.600 | - | - | -3.00 ± 7.92 | -3.00 ± 7.68 | - | - |
| 5 | 11.000 ± 9.250 | 33.000 ± 12.750 | - | - | 5.50 ± 8.25 | -0.50 ± 20.00 | - | - |
| 6 | 27.000 ± 7.600 | 30.000 ± 8.960 | - | - | 0.00 ± 4.32 | 17.00 ± 12.16 | - | - |
| 7 | 8.000 ± 5.556 | 7.000 ± 10.518 | <.001 | <.001 | 2.00 ± 5.31 | 3.00 ± 11.71 | <.001 | <.001 |
| 8 | 5.000 ± 3.500 | 4.500 ± 7.750 | - | - | 2.00 ± 1.75 | 0.00 ± 3.50 | - | - |
| 9 | 4.000 ± 4.667 | 0.000 ± 1.111 | .008 | .125 | 2.00 ± 8.96 | 0.00 ± 2.86 | .063 | .625 |
| 10 | 7.000 ± 3.630 | 0.000 ± 1.407 | .008 | 0.500 | 0.00 ± 2.47 | 0.00 ± 1.04 | .500 | .500 |
| 11 | 0.000 ± 2.519 | 0.000 ± 2.815 | .148 | .900 | .00 ± 1.91 | 0.00 ± 4.38 | .900 | .628 |
| 12 | 2.500 ± 3.830 | 8.000 ± 5.620 | .010 | .001 | 1.50 ± 6.20 | 2.00 ± 8.30 | .395 | .243 |
| 13 | 0.000 ± 1.045 | 1.000 ± 1.619 | .426 | .008 | 0.00 ± 1.86 | -1.00 ± 4.03 | .383 | .209 |
| 14 | 2.750 ± 2.061 | 4.250 ± 4.235 | .001 | .001 | 0.00 ± 1.33 | 0.50 ± 2.06 | .510 | .982 |
| 15 | 3.000 ± 4.098 | 1.000 ± 3.467 | .003 | .005 | 2.00 ± 3.25 | 0.00 ± 4.69 | .168 | .625 |
| 16 | 3.000 ± 2.441 | 1.750 ± 3.477 | .001 | .044 | -0.50 ± 1.94 | 0.00 ± 1.97 | .291 | .313 |
| 17 | 2.750 ± 2.929 | 1.000 ± 3.806 | .010 | .031 | -1.00 ± 2.41 | 0.00 ± 3.14 | .191 | .573 |

The IDs per row refer to the numeric identifiers of treatment conditions listed in Table 1.

**Figure 2.**
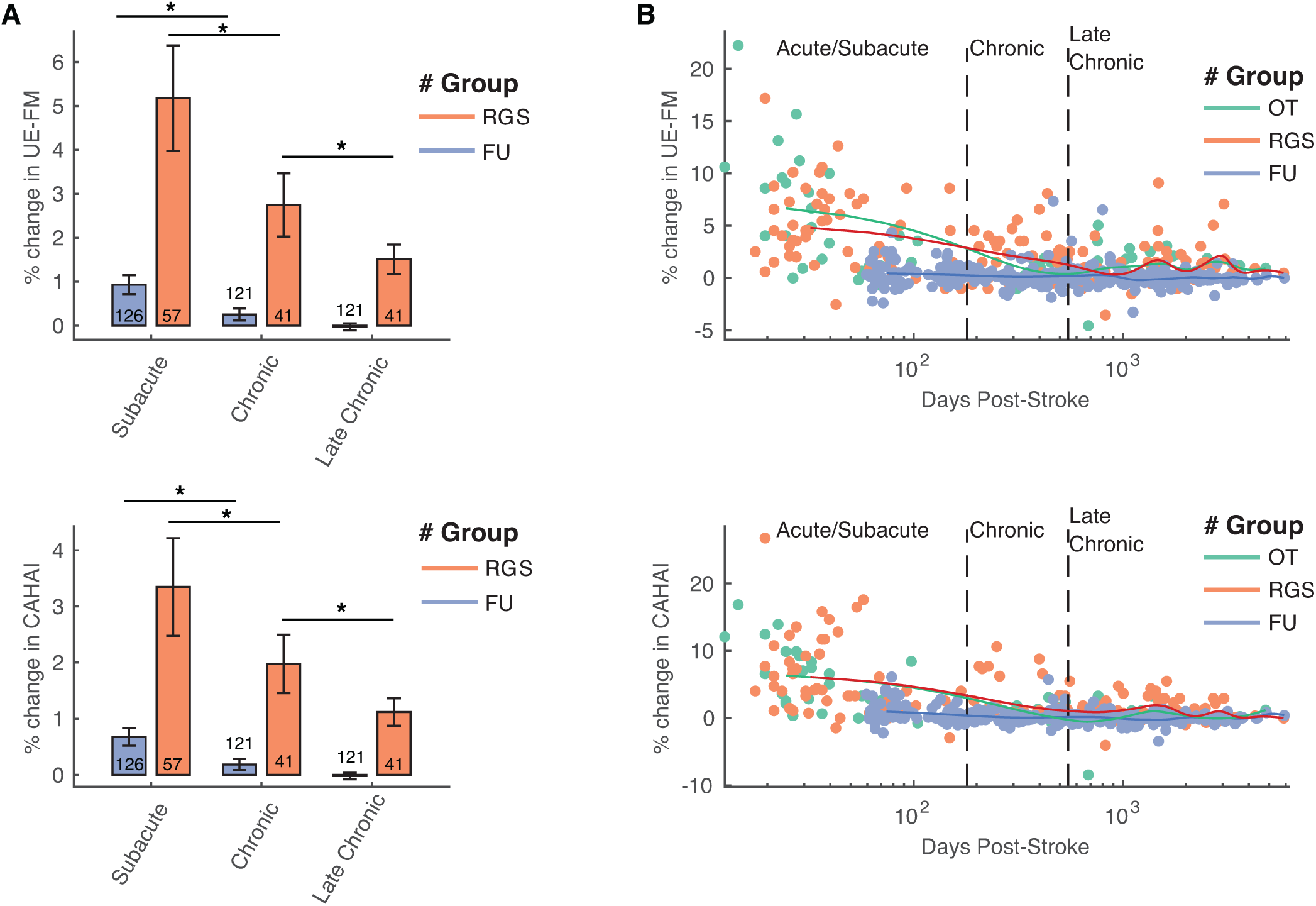
A. Averaged normalized improvement rates per week of the RGS group and Follow-up (FU) for UE-FM (top) and CAHAI scales (bottom), by patient’s chronicity at the time of the evaluation. The number of observations is indicated within or above each bar. B. Comparison of the RGS, Occupational Therapy (OT) and Follow-up (FU) measures of normalized improvement rates per week for UE-FM (top) and CAHAI (bottom) scales, by patient’s chronicity at the time of the evaluation. Solid lines indicate the estimated averages. Vertical dashed lines indicate the limits of the three chronicity categories.

### Revealing an extended critical window of recovery

In order to combine the impact of all the studies and to further examine the effects of chronicity on the recovery potential metric (see Eq. 1 in Methods), we apply a bootstrapping method that merges clinical evaluations from all the samples. Clustering the data in three chronicity categories, we observe a 1 to 2 % normalized improvement (see Equation 1 in Methods) on the UE-FM scale per week of training, dependent on the patient’s chronicity, with a mean improvement of 5.2 ± 1.0 SD % in acute/subacute, and 2.7 ± 0.6 SD % and 1.4 ± 0.3 SD % in early and late chronic patients respectively (Figure 2 A). We observe no significant differences in recovery rates between Occupational Therapy and RGS groups. The change on the CAHAI scale is in the 1 to 4 % range of normalized recovery per week of training with a mean improvement of 5.2 ± 1.0 SD % in acute/subacute, and 2.7 ± 0.6 SD % and 1.4 ± 0.3 SD % in early and late chronic patients respectively. We found statistically significant differences between acute/subacute and early chronic for all groups, even at follow-up (p< 0.01, Wilcoxon Rank-Sum). However, only in the RGS group the early chronic group showed higher recovery rates than the late chronic group (p< 0.05, Wilcoxon Rank-Sum), revealing a long-lasting gradient of recovery potential that remains visible across the first 18 months post-stroke (Figure 2 B). This effect was not present in the Follow-up measures. Due to the low sample size in the Occupational Therapy group we could not apply the bootstrapping and no gradient was detected among this patients. The gradient found in the RGS group could not be explained by the patient’s age (Spearman’s correlation r< 0.003, p> 0.96) and neither by the patient’s baseline impairment score (Spearman’s correlation r< 0.052, p> 0.43 for FM; < 0.006, p> 0.93 for CAHAI). Notice that the design of this retrospective analysis controls for additional confounding variables, since all patients included in this analysis were recruited according to common inclusion and exclusion criteria with respect to age, motor impairment severity, cognitive impairment severity, type of stroke (Oxford Classification), hand dominance, absence of a second stroke, and gender. We emphasize that in the dataset used for our analysis, none of these variables correlate with the patients’ chronicity and therefore none of them can explain the uncovered gradient.

## DISCUSSION

In opposition to motor learning and compensation, true recovery has been conceptualized as the biological process underlying a change in UE-FM scores from baseline in a thirty days time-window (Bernhardt et al. 2017). Here we show that improvements on the UE-FM scale, related to true recovery, can also occur at later stages post-stroke (e.g. even 18 months post-stroke), however capturing this effect may require large homogeneous datasets and analytical methods with enhanced accuracy such as a bootstrapping technique. By analyzing clinical recovery scores from stroke patients with variable chronicity but homogeneous motor and cognitive impairment levels, we are able to detect a gradient of recovery potential that extends beyond 18 months post-stroke. In line with the previous literature, our data illustrates the correlation of *δ* UE-FM and 5 CAHAI scores at acute stages (Beebe and Lang, 2009; Rabadi and Rabadi, 2006), while at chronic stages these scales dissociate, possibly due to the introduction of compensatory mechanisms (Rabadi and Rabadi, 2006). Furthermore, in all studies analyzed, the obtained gains due to rehabilitation were retained at follow-up (Supplementary Information, Figure 3).

The results presented in this study do require further investigation for a number of reasons. First, the observation of non-significant effects in one of the nine studies may be due to low statistical power and risk of bias, however since this study contributes to the final analysis with only 18 data points, it seems highly unprovable that it may have biased our results. Second, the functional relevance of the detected improvements is marginal at chronic stages. The UE-FM minimal clinically important difference (MCID) has been defined as a constant value across the patient’s chronicity spectrum, however, this assumption lacks support since, according to the gradient of recovery potential revealed by our analysis, one would expect chronicity-dependent variability in the UE-FM’s measurement error and the patient’s perceived improvement thresholds. Thus, our findings suggest a need for redefining MCID thresholds. Third, the uncovered gradient of sensitivity to therapy may not be specific to VR-based interventions. Due to the low number of early chronic patients in the Occupational Therapy group, we could not perform a bootstrapping analysis in this group and we were not able to explore the related recovery dynamics. Future studies should explore the influence of conventional therapy and alternative rehabilitation approaches on different chronicity windows.

## 4 CONCLUSION

Using a VR-based rehabilitation protocol and a novel bootstrap analysis for unifying the results from eleven VR-based interventions at different stages post-stroke we observed structural and functional improvement over the treatment period at all stages post-stroke. This effect displayed a gradient of sensitivity to treatment that faded out exponentially and reached asymptotic levels after one year and a half post-stroke. These findings call for an urgent scientific effort to redefine the ‘critical window of recovery’ and highlight the need for providing therapy to patients at the chronic and late chronic stage.

## AUTHOR CONTRIBUTIONS

Conception and design of the study: BRB, AD, MM, MC, SB, EDO, AC, SR, and PV; implementation: BRB, AD, MM; data acquisition: BRB, MM, MC, SB; analysis and interpretation: BRB, AD, MM, MC, SB, EDO, AC, SR, and PV; and manuscript preparation: BRB, AD, MM, MC, SB, EDO, AC, SR, and PV.

## ACKNOWLEDGMENTS

The authors gratefully acknowledge the participation of the patients and thank Irene Camacho and Estefanía Montiel for their assistance in the collection of the data.

## DATA AVAILABILITY STATEMENT

The complete dataset and analysis source-code will be open and freely available on publication. This resource will contain the following datasets:

**Dataset D1.** “PatientsDemographicsAndClinicalScreening.xlxl”: Demographical data and clinical screening information (age, gender, chronicity, center ID, stroke type, oxford classification, affected arm, arm dominance, presence of aphasia, days after stroke).

**Dataset D2.** “ClinicalScalesAll.csv”: Recovery scores from 219 hemiparetic stroke patients evaluated using the Upper Extremity section of the Fugl-Meyer, CAHAI, and Barthel clinical scales at multiple time points (baseline, end of the treatment, and follow-up periods).

## Revealing an extended critical window of recovery post-stroke

4.0.1 Figures

**Figure S1.**
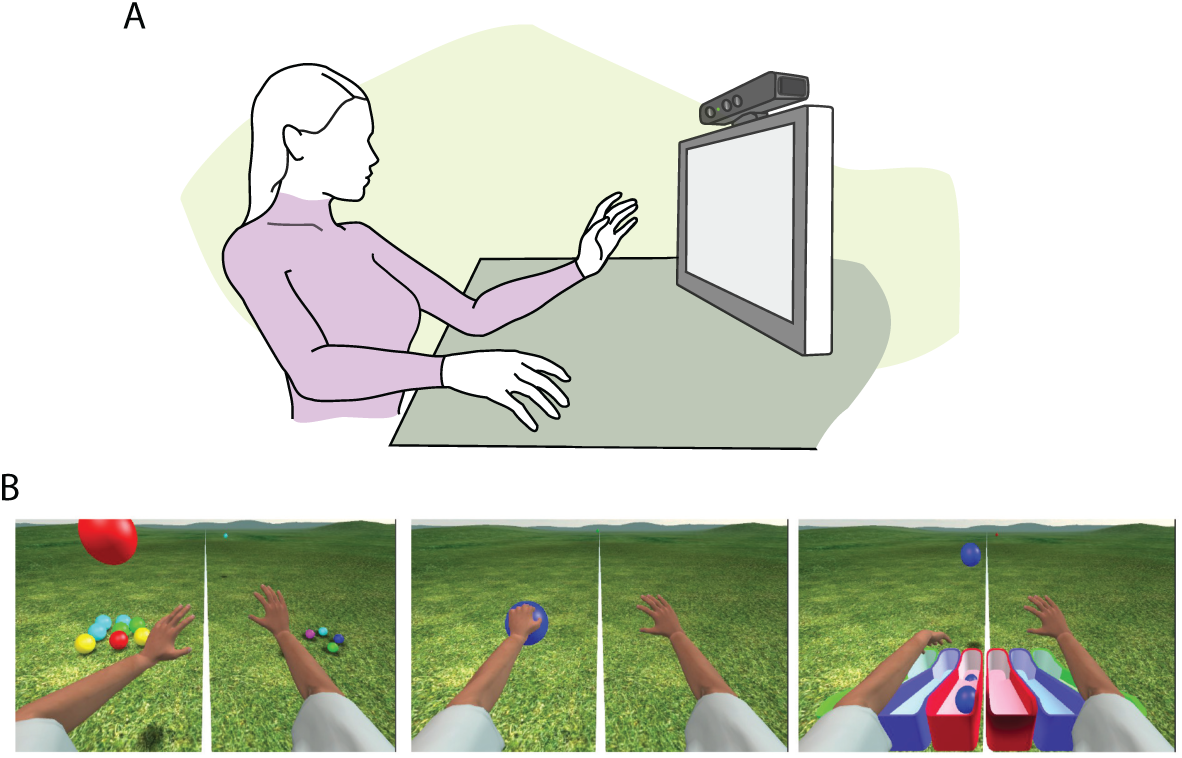
Illustration of the Rehabilitation Gaming System (RGS). A: The system consists of a PC, a 17 inch LCD display, a vision-based motion capture device (Kinect 360, Microsoft, Seatle USA) positioned on top of the screen. The virtual tasks logic and graphics were implemented using the Unity 3D (Unity Technologies, San Francisco, USA) and Torque (Garage Games, Las Vegas, NV, USA) computer game engines. The vision-based motion tracking device and data gloves capture the joint movements of the user’s torso, shoulders, elbows, and fingers, and map them onto an avatar through a biomechanical model, thus mimicking the movements of the user. Arm and finger movements are displayed on a screen in a first-person perspective, realizing a paradigm that combines goal-oriented action execution, motor imagery, and action observation. B: the Spheroids task follows a proximal to distal training progression where the users are asked to intercept spheres that move towards them (left). Followed by grasping (middle), and placing the spheroids in color matched fashion (right). The scenario is adapted to the performance of the user by controlling the difficulty of the task as defined by the frequency, the speed and the horizontal range of the spheres.

**Figure S2.**
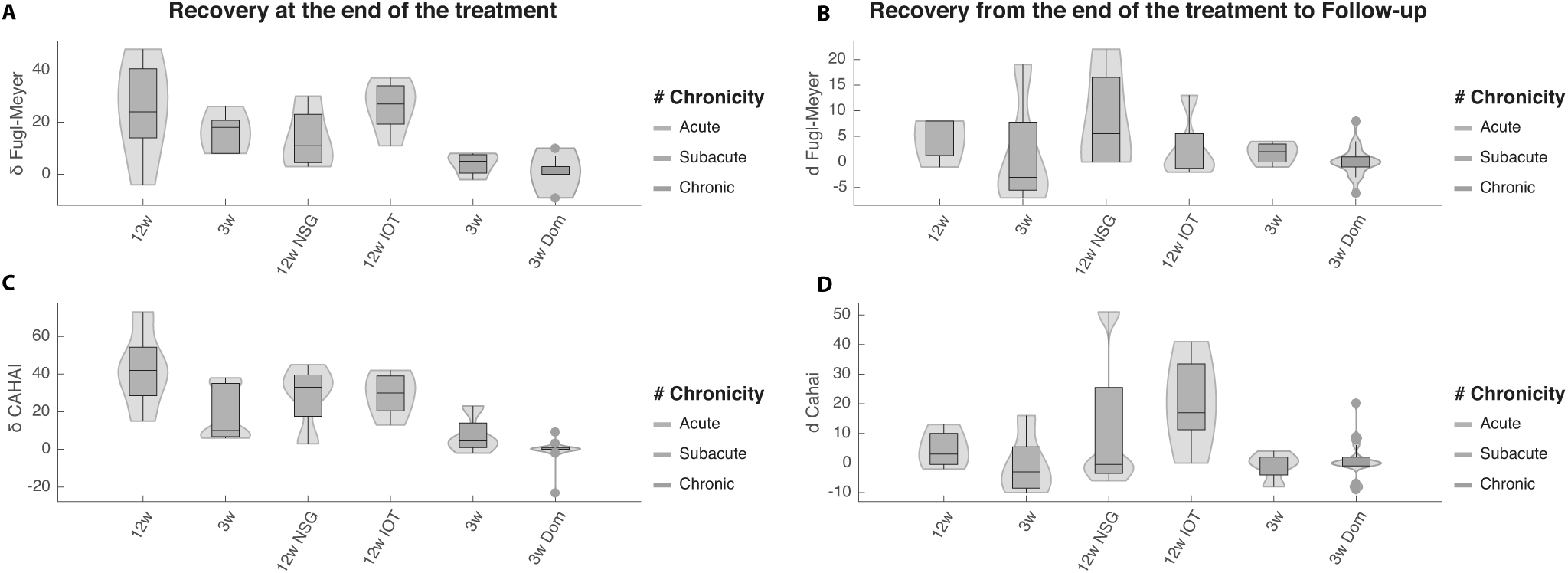
Improvement Rate per Week for UE-FM (A-B) and CAHAI (C-D), by patient’s chronicity at the time of the evaluation. Effect of RGS-based training at the end of the treatment (A and C), and during the follow-up (B and C). The length of each intervention is indicated in weeks (w). Notice that the horizontal axis refers to the studies listed in Table 2 and follow the same order. Shaded areas indicate the data distribution color coded according to the chronicity of stroke patients participating in each study: acute (green), subacute (orange), and early chronic stage (blue).

**Figure S3.**
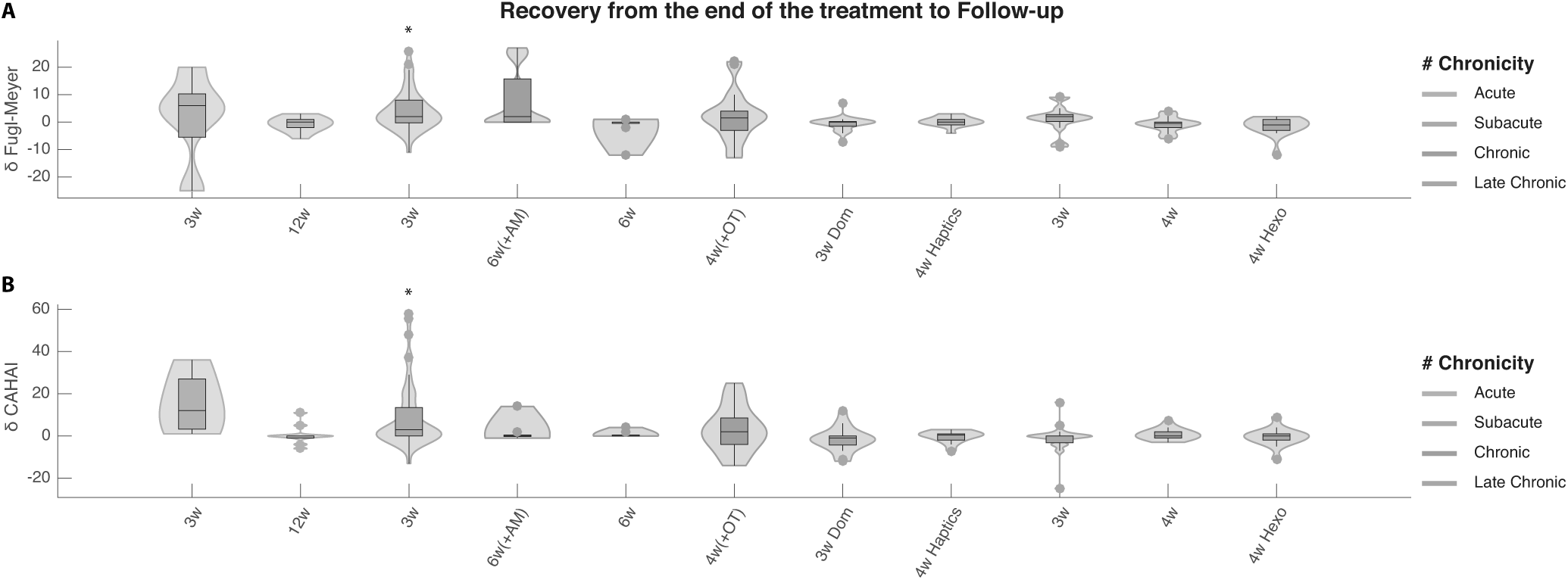
Effect of RGS-based treatment from the end of the therapy to the end of the follow-up period. Impact measured on upper limb motor function (UE-FM) (Top) and performance in iADLs (CAHAI) (Bottom). The effect represents a change in each scale from the end of treatment to follow-up evaluation. The length of each intervention is indicated in weeks (w). Notice that the horizontal axis refers to the RGS conditions listed in Table 2 and follow the same order. Shaded areas indicate the data distribution color coded according to the chronicity of stroke patients participating in each study: acute (green), subacute (orange), and early (blue) to late (purple) chronic stage. * for p-value< 0.05 and ** for p-value¡ 0.01.

**Table S1:**
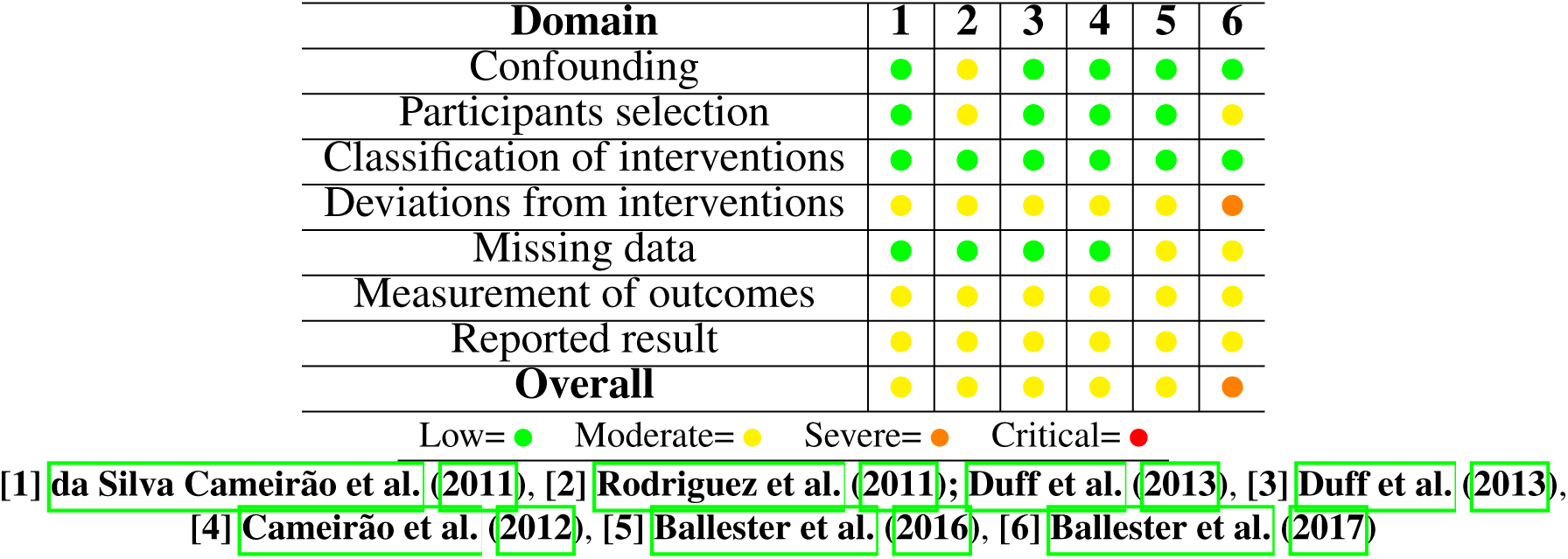
Risk Of Bias. Illustration of different Risk Of Bias In Non-randomized Studies of Interventions judgments (ROBINS-I) for each study included (Ballester et al., 2016, 2017; da Silva Cameirão et al., 2011; Cameirão et al., 2012; Duff et al., 2013; Rodriguez et al., 2011).

